# Non-canonical autophagy drives alternative ATG8 conjugation to phosphatidylserine

**DOI:** 10.1101/2020.05.14.096115

**Authors:** Joanne Durgan, Alf H. Lystad, Katherine Sloan, Sven R. Carlsson, Michael I. Wilson, Elena Marcassa, Rachel Ulferts, Judith Webster, Andrea F. Lopez-Clavijo, Michael J. Wakelam, Rupert Beale, Anne Simonsen, David Oxley, Oliver Florey

## Abstract

Autophagy is a fundamental catabolic process essential for development, homeostasis and proper immune function ^1^. During autophagy, a cascade of ATG proteins target intracellular cargoes for lysosomal degradation and recycling ^2^. This pathway utilises a unique post-translational modification, the conjugation of ATG8 proteins to phosphatidylethanolamine (PE) at autophagosomes, which modulates cargo selection and maturation. ATG8 lipidation also occurs during non-canonical autophagy, a parallel pathway involving Single Membrane ATG8 Conjugation (SMAC) to endolysosomal compartments, which plays a key role in phagocytosis and other processes ^3^. It has been widely assumed that SMAC involves the same lipidation of ATG8 to PE, but this has yet to be formally tested. Here, we show that ATG8 undergoes alternative lipidation to phosphatidylserine (PS) during non-canonical autophagy/SMAC. Using mass spectrometry, we find that activation of SMAC, by pharmacological agents ^4,5^, or during non-canonical autophagy processes such as LC3-associated phagocytosis ^6,7^ and Influenza A virus infection ^8^, induces the covalent conjugation of ATG8 to PS, as well as PE. This alternative lipidation event is dependent on the ATG16L1 WD40 domain, and occurs at PS enriched endolysosomal membranes. Importantly, we find that the ATG8-PS and ATG8-PE adducts are differentially delipidated by isoforms of the ATG4 family, indicating significant molecular distinctions and mechanisms between these two species.

Together, these results provide an important new insight into autophagy signalling, revealing an alternative form of the hallmark ATG8-lipidation event, so widely used to define and assay autophagy. Furthermore, ATG8-PS lipidation provides a specific ‘molecular signature’ for non-canonical autophagy, uncovering a novel means of detecting and monitoring this emerging pathway.

## Main

A defining feature of autophagy is the lipidation of ATG8, a family of ubiquitin-like proteins including mammalian LC3s (A/B/B2/C) and GABARAPs (GABARAP/L1/L2)^9^. Nascent pro-ATG8 is first primed by a cysteine protease, ATG4, to expose a conserved aromatic-Gly motif at its C-terminus ^10^. A ubiquitin-like conjugation system, comprised of ATG7 (E1-like), ATG3 (E2-like) and ATG16L1/ATG12/ATG5 (E3-like), then drives the covalent ligation of this glycine to a lipid, phosphatidylethanolamine (PE), via an amide bond to its headgroup (Extended Data Fig. 1a) ^11,12^. This is a unique post-translational modification that recruits ATG8 to autophagosomal membranes, where it plays an important role in cargo loading and autophagosome maturation ^9,13^. The associated relocalisation of ATG8s, and the characteristic protein bandshift between the unlipidated (ATG8-I) and lipidated (ATG8-II) forms, are widely used to define and assay autophagy-related processes^14,15^.

A second phospholipid, phosphatidylserine (PS), also bears an amino group in its head moiety (Extended Data Fig. 1b), which can be conjugated to ATG8 *in vitro* ^16^. However, *in vivo*, ATG8 lipidation is reported to occur exclusively to PE, in both yeast ^11^ and mammalian cells ^16^. The mechanism underlying cellular specificity is not fully understood, but physiological pH and phospholipid composition may prohibit alternative lipidation to PS ^17,18^. Alternatively, ATG8 conjugated to PS may be subject to rapid delipidation.

It is increasingly clear that the autophagy machinery mediates critical, parallel functions in other vital cellular processes ^3^. During non-canonical autophagy, a subset of core ATG proteins (including ATG7/3/12/5/16L1), but not the upstream regulators (FIP200/ULK/ATG13), target various endolysosomal compartments for <u>S</u>ingle <u>M</u>embrane <u>A</u>TG8 <u>C</u>onjugation (SMAC). LC3-associated phagocytosis (LAP) is an important example of this pathway, where LC3 conjugation to phagosomes, housing pathogens or apoptotic debris, plays a key role in modulating the immune response ^7^, inflammation ^6,19,20^, antigen presentation ^8,21^, vision ^22^ and tumour cell tolerance ^23^. SMAC is also active during macropinocytosis, entosis ^24^ and LC3-associated endocytosis (LANDO) ^25^. In each case, the non-canonical autophagy/SMAC pathway culminates in ATG8 lipidation, which has been widely assumed to represent PE conjugation ^6,23,24^. However, the identity of the modified-ATG8 has not been formally tested in this context.

## Mass spectrometric analysis of ATG8 lipidation

To investigate the nature of ATG8 lipidation during SMAC, we took a mass spectrometric approach. GFP-tagged ATG8 proteins were expressed in cells of different genetic backgrounds, and treated with different pharmacological stimuli, to drive ATG8-lipidation associated with either canonical autophagy, or SMAC (Fig. 1). GFP-ATG8 was then immunoprecipitated, base treated to remove phospholipid acyl chains (leaving only the headgroup conjugated), and subjected to proteolytic cleavage with AspN protease (Extended Data Fig. 1c). The resulting ATG8 C-terminal peptides, in their unmodified form, or covalently conjugated to a phospholipid headgroup, were analysed by liquid chromatography-tandem mass spectrometry (LC-MS/MS). Where linked to glycerophosphoethanolamine (from PE), this peptide has a mass of 1923.7996; if conjugated to glycerophosphoserine (from PS), the expected mass would be 1967.7894.

**Figure 1:**
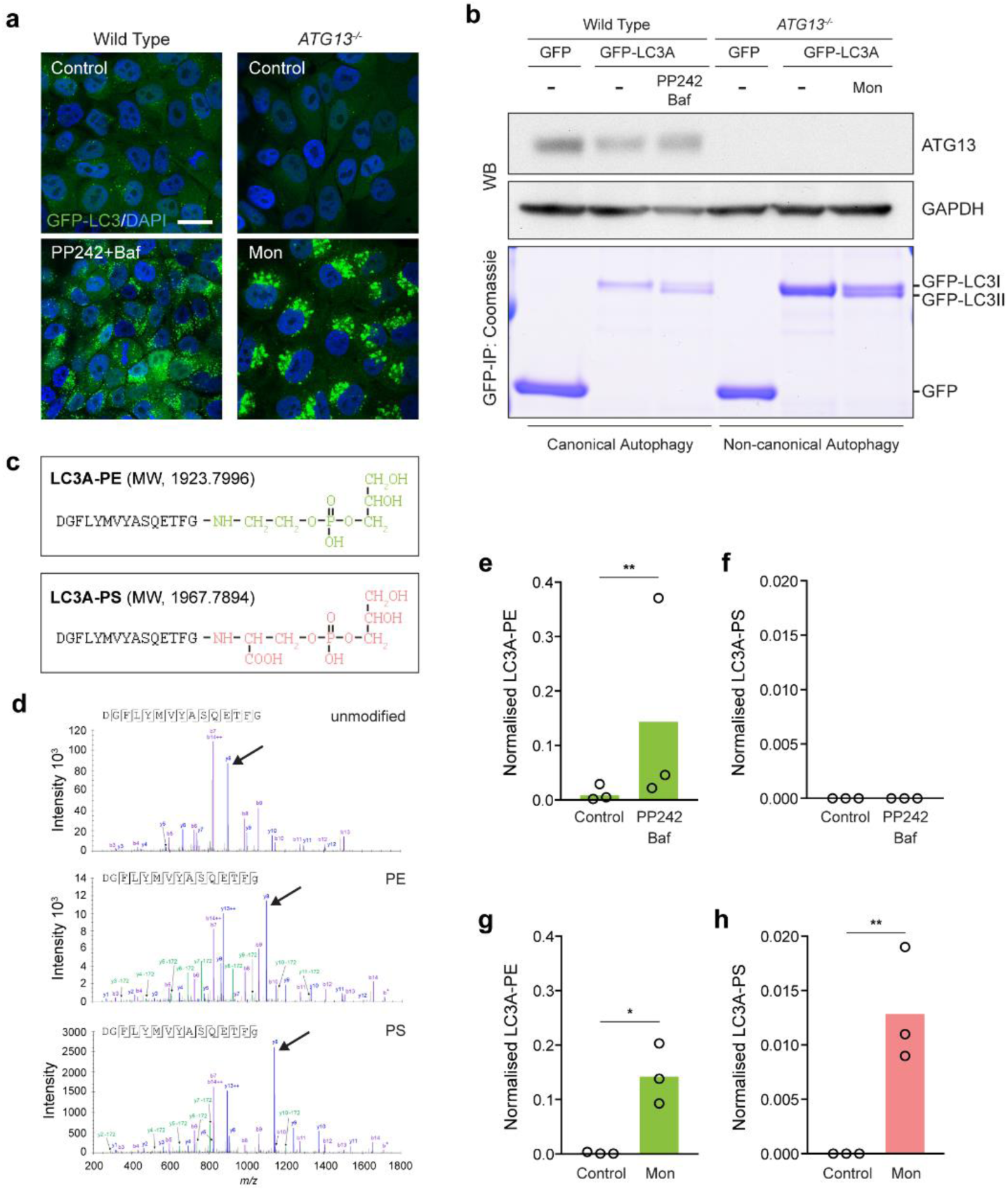
Pharmacological activation of non-canonical autophagy promotes ATG8-PS lipidation in cells. Wild type MCF10A GFP-LC3A cells were treated with 1 μM PP242/100 nM BafA1 for 60 mins. *ATG13*^*-/-*^ cells were treated with 100 μM monensin (Mon) for 40mins. **a**, Confocal images of GFP-LC3A and DAPI. Scale bar: 20 μm. **b**, Western blotting for ATG13 or GAPDH, GFP-IPs were visualised by Coomassie staining. **c**, C-terminal peptides of LC3A conjugated to either the PE or PS headgroup. Predicted MWs are indicated. **d**, Representative CID mass spectra of unmodified, PE or PS modified LC3A peptides. y-ions (C-terminal) undergo monoisotopic mass shift upon modification: 197.05, glycerophosphoethanolamine (from PE); 241.04, glycerophosphoserine (from PS); y8 peaks are highlighted (arrow heads) as examples. Some y-ions give a secondary fragment consistent with neutral loss of phosphoglycerol (172). As expected, b-ions (N-terminal) do not shift. b14 is characteristically absent from the unmodified peptide, but can be observed in the modified peptides, along with b* (cleavage between Gly and the head group), confirming the C-terminal amide linked modification. **e-h**, Normalized mass spectrometry analysis of LC3-PE and LC3A-PS in WT **e, f**, and *ATG13*^*-/-*^ **g, h**, cells. Data represent means from 3 independent experiments, **p<0.002, *p<0.004, ratio paired t-test.

As proof-of-concept, ATG8 lipidation associated with canonical autophagy was induced in wild type cells expressing GFP-LC3A, by co-treatment with an mTOR inhibitor (PP242) and a V-ATPase inhibitor (Bafilomycin A1, BafA1), which induce and accumulate autophagosomes respectively. As expected, GFP-LC3A relocalises to punctate autophagosomes upon PP242/BafA1 treatment (Fig. 1a) and a faster migrating, lipidated band is observed by coomassie staining (GFP-LC3II, Fig. 1b). By mass spectrometry, this lipidation event corresponds exclusively to the covalent conjugation of PE (Fig. 1c, d, e), with no detectable trace of PS (Fig. 1f). These findings are consistent with published work, in which activation of autophagy *in vivo* induces the selective conjugation of ATG8 to PE in both yeast and mammalian cells^11,16^.

To investigate ATG8 lipidation during non-canonical autophagy, *ATG13*^-/-^ cells, deficient in canonical autophagy, were treated with monensin, a known activator of SMAC ^5,26,27^ (Fig. 1a, b). Consistent with previous work ^4,5,8^, these conditions yield specific activation of SMAC, inducing GFP-LC3A recruitment to endolysosomal membranes, and a lipidation-associated bandshift (Fig. 1a, b), with no significant effect on global lipid composition (Extended Data Fig.2). Strikingly, under these conditions, mass spectrometry detects LC3A conjugated to both PE and PS (Fig. 1c, d, g, h). These data provide the first evidence for *in vivo* cellular LC3-PS conjugation. To extend these analyses, additional ATG8 isoforms were examined (Extended Data Fig. 3). Interestingly, monensin also drives the PS-lipidation of GABARAP isoforms, confirming that PS-lipidation is conserved across multiple ATG8 family members. Using normalised peak areas to estimate relative abundance, ATG8-PS represents approximately 10% (LC3A) to 30% (GABARAP) of the lipidated form, under these conditions.

Collectively, these data establish that ATG8-PS lipidation can occur in cells, and indicate that non-canonical autophagy/SMAC drives this distinctive modification.

## ATG8-PS lipidation during physiological non-canonical autophagy/SMAC

To extend these findings to more physiological processes, SMAC was first analysed in the context of LC3-associated phagocytosis (LAP). Using the J774A.1 macrophage model, GFP-LC3B recruitment to phagosomes housing IgG-coated beads was analysed in the presence or absence of BafA1 (Fig. 2a, b). BafA1 inhibits SMAC, which is V-ATPase dependent ^4,28^, in contrast to its effects on canonical autophagy (noted in Fig. 1a, b). As expected for LAP, BafA1 does not inhibit the number of phagosomes formed (Fig. 2c), but does reduce levels of lipidated GFP-LC3B-II (Fig. 2d). These data also verify that the majority of enriched GFP-LC3B derives from phagosomes, not contaminating autophagosomes (where BafA1 would instead increase GFP-LC3BII due to block of autophagosome flux). Importantly, using this system, we demonstrate that induction of LAP drives the alternative lipidation of LC3 with PS, as well as with PE (Fig. 2 e, f). In this case, LC3B-PS accounts for approximately 25% of the lipidated species, and is reduced by BafA1 even more robustly than LC3-PE.

**Figure 2:**
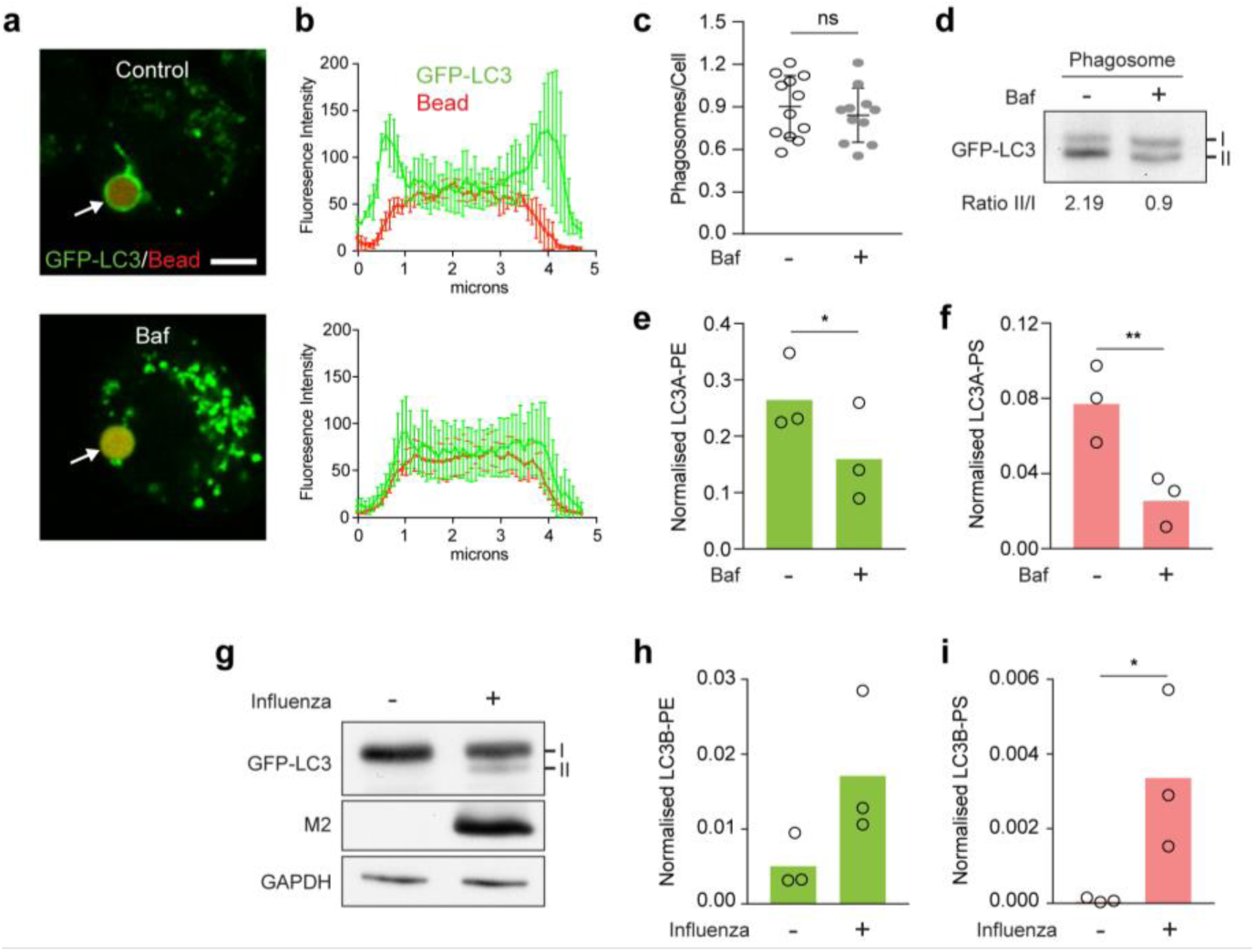
LC3-PS lipidation occurs during LC3-associated phagocytosis (LAP) and Influenza A Virus (IAV) infection. **a**, J774A.1 macrophage treated with IgG coated beads to induce LAP. Representative confocal images show GFP-LC3A localisation -/+ BafA1 (100 nM). Scale bar: 5 μm **b**, Signal intensity profile of phagosomal GFP-LC3. Data represent mean ±S.D. from 3 phagosomes. **c**, Quantification of phagocytosis ±S.D. >10 fields of view. **d**, Representative western blot of GFP-LC3A from phagosome fraction with ratio of LC3II/LC3I. **e, f**, Normalised mass spectrometry analysis of LC3A-PE and LC3A-PS from phagosome fractions. **g**, HCT116 cells infected with Influenza A Virus (IAV) PR8 and analysed by western blot for GFP-LC3B, viral M2 and GAPDH. **h, i** Normalised mass spectrometry analysis of LC3B-PE and LC3A-PS -/+ IAV infection. Data represent means from 3 independent experiments, **p<0.008, *p<0.02, paired t test.

To investigate an additional physiological trigger, Influenza A Virus (IAV) infection was assessed, in which the viral M2 proton channel drives SMAC ^8^. Upon infection and M2 protein induction, GFP-LC3B was relocalised (Extended Data Fig. 4) and lipidated (Fig. 2g). Under these conditions, mass spectrometric analysis detects GFP-LC3B conjugation to both PE and PS, with PS representing around 20% of the total lipidated species (Fig. 2h, i).

Together, these data establish that ATG8-PS lipidation occurs broadly upon induction of SMAC via pharmacological activation, LAP or Influenza A Virus infection.

## Molecular mechanisms of ATG8-PS lipidation

To define the molecular mechanisms underpinning differential ATG8 lipidation, the contribution of ATG16L1 was examined. ATG16L1 acts as a molecular hub, coordinating autophagy related pathways, via distinct domains that support either canonical or non-canonical signalling ^8,26,29,30^. We previously identified key residues within the ATG16L1 WD40 domain which, when mutated (e.g. K490A), render cells competent for canonical autophagy, but deficient for SMAC ^8,26^, and can thus be used to dissect these pathways. A panel of *ATG16L1*^*-/-*^ HCT116 cells, reconstituted with either wild type (WT) ATG16L1 or the K490A mutant, were thus analysed (Fig. 3a). In WT cells, GFP-LC3B relocalises and is lipidated to both PE and PS upon monensin treatment (Fig. 3b-d). In K490A mutants, a small but reproducible increase in LC3-PE is detected upon monensin treatment, corresponding to a block of basal autophagy flux, and these cells support robust LC3-PE lipidation upon PP242/BafA1 treatment (Extended Data Fig. 5). In contrast, K490A mutant cells are completely deficient in SMAC associated LC3-PS lipidation. Similarly, LC3-PS can be detected in WT, but not K490A, RAW267.4 macrophage undergoing LAP (Extended Data Fig. 6).

**Figure 3:**
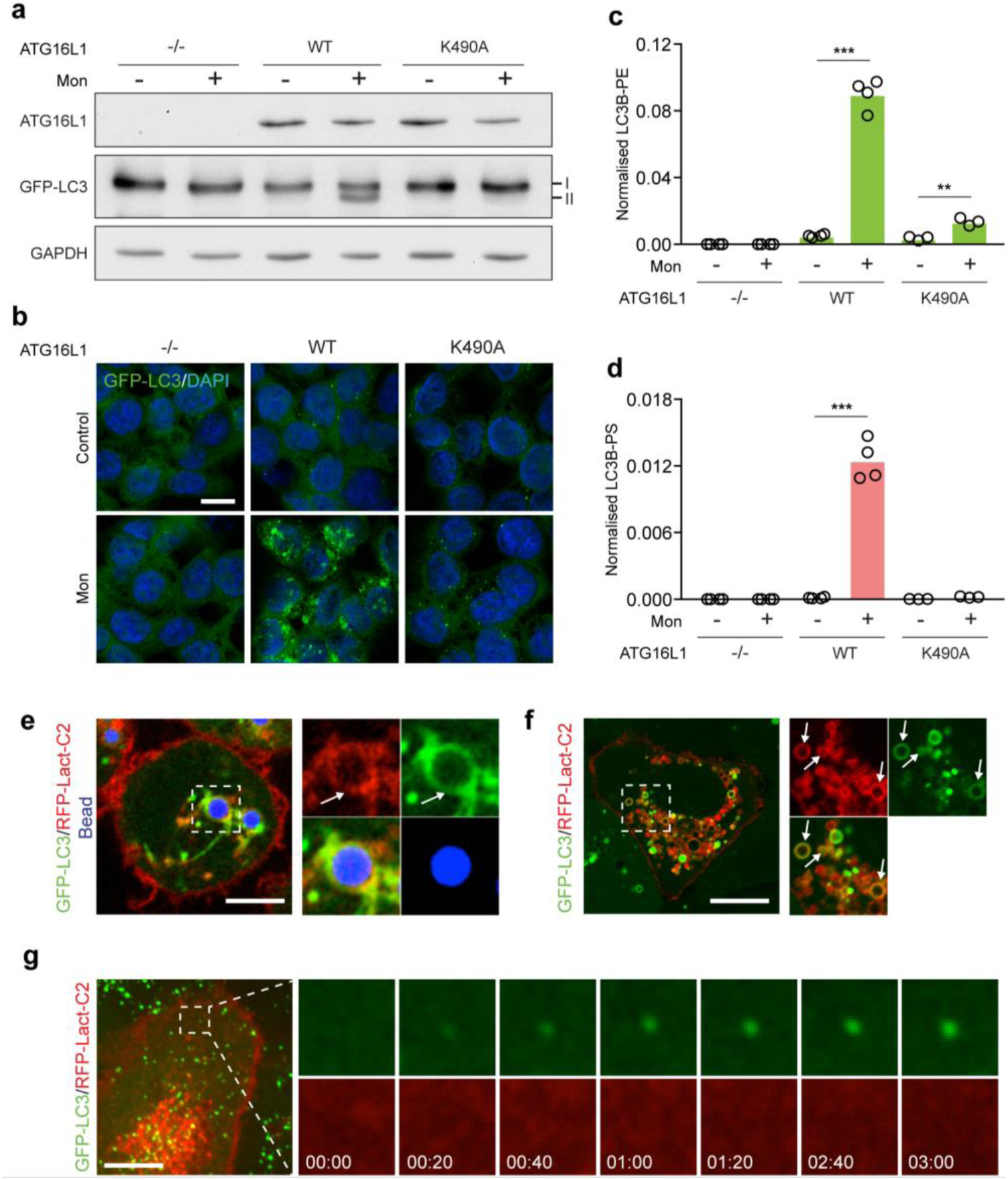
The ATG16L1 WD40 domain supports alternative LC3 lipidation, at PS enriched membranes. HCT116 *ATG16L1*^*-/-*^ cells, re-expressing WT or K490A ATG16L1, and GFP-LC3B, were treated -/+ 100 μM monensin (Mon) for 40 mins. Cells were analysed by **a**, western blot, **b**, confocal microscopy (scale bar: 20 μm) and **c, d**, Mass spectrometry for normalised LC3B-PE or PS. Data represent 3-4 independent experiments, ***p<0.0003, **p<0.006, paired t test. **e-g**, Confocal images of **e**, GFP-LC3A and RFP-Lact-C2 (PS sensor) in J774.A1 cells during LAP of IgG-beads. Scale bar: 5 μm, arrows denote a GFP-LC3 positive phagosome, **f**, Monensin treated MCF10A; scale bar: 10 μm, arrows denote GFP-LC3A positive lysosomes. **g**, Live confocal imaging of PP242 treated MCF10As. Scale bar: 5 μm. Cropped time-lapse frames, min:sec.

Together, these data indicate that ATG8-PS lipidation is completely dependent on the machinery of non-canonical autophagy/SMAC.

ATG16L1, in complex with ATG5 and ATG12, serves to direct the site of ATG8 lipidation ^31^. We thus reasoned that alternative ATG8 lipidation may result, at least in part, from differences in lipid composition at the distinct membranes targeted by ATG16L1 during SMAC versus autophagy ^32^. To investigate this, a fluorescent sensor for PS (Lact-C2) ^33^ was expressed in cells undergoing different autophagy related processes. Notably, PS is enriched at various LC3-positive compartments during SMAC, including phagosomes and lysosomes (Fig. 3e, f), as well as macropinosomes and entotic corpse vacuoles (Extended Data Fig. 7). In contrast, PS could not be detected on forming autophagosomes (Fig. 3g). These data support a simple model in which local PS availability may influence the identity of LC3 lipidation, although other regulatory mechanisms may also operate.

Collectively, these data indicate that the molecular machinery of non-canonical autophagy/SMAC, such as the ATG16L1 WD40 domain, direct ATG8-PS conjugation at PS enriched endolysosomal single membranes.

## Differential PE/PS delipidation by ATG4s

Finally, the molecular consequences of differential ATG8 conjugation were investigated, with a focus on ATG4, the dual activity proteases that regulate ATG8s. ATG4s first prime pro-ATG8s for lipidation by proteolytic cleavage, then subsequently catalyse their delipidation. To explore whether ATG4 processing may be sensitive to alternative ATG8 lipidation, the conjugation of either PE or PS was computationally modelled onto the LC3(120)-ATG4B co-complex structure (Fig. 4a) ^34^. These phospholipids differ by just a single carboxyl group, which confers extra bulk and negative charge to PS. Modelling suggests that this distinctive carboxyl moiety would juxtapose with ATG4B Trp142, a residue critical for structure and activity ^35^. As such, the additional PS carboxyl group may limit freedom of movement and sterically hinder its delipidation. To test this experimentally, a mixed pool of SMAC-induced LC3A-PE and LC3A-PS was enriched from cells and incubated with recombinant ATG4B *in vitro* (Fig. 4b). Strikingly, while LC3-PE undergoes robust delipidation through time (Fig. 4c), and across replicate experiments (Fig.4d), LC3-PS is largely resistant to deconjugation under these conditions. These findings confirm that differential LC3 lipidation influences ATG4B-mediated deconjugation, revealing an important functional outcome for this alternative modification. Interestingly, these findings are consistent with a previous *in vitro* study of GABARAPL1 conjugated liposomes, delipidated by ATG4A, B or C ^36^, indicating that a reduced rate of ATG8-PS cleavage is likely shared among multiple isoforms of both ATG8 and ATG4.

**Figure 4:**
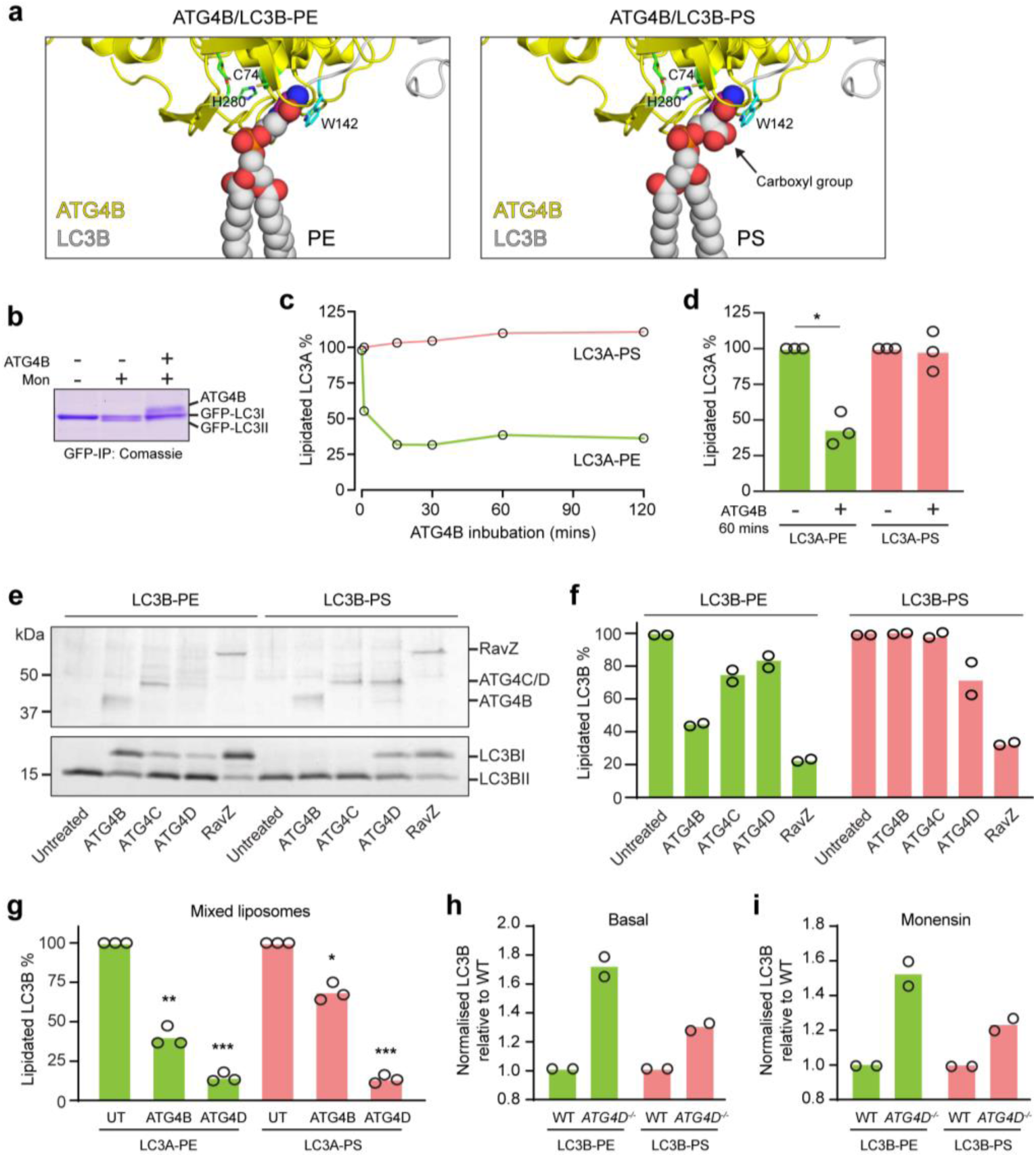
LC3A-PS and LC3A-PE undergo differential delipidation by the ATG4 family. **a**, Molecular modelling of LC3-PE and LC3-PS in complex with ATG4B (based on 2Z0D.pdb), with critical ATG4B catalytic residues marked. **b**, Coomassie staining of GFP-LC3A immunoprecipitated from monensin (mon) treated MCF10A *ATG13*^*-/-*^ cells, then incubated -/+ recombinant ATG4B for 60 mins. **c**, Mass spectrometry analysis of LC3-PE and LC3-PS from monensin treated cells following a time course of ATG4B incubation and **d**, 3 independent experiments at 60 mins. Data represent means normalised to time 0, *p<0.01, paired t test. **e-i**, Liposome-based delipidation assays with purified ATG4s and RavZ. **e**, GFP-LC3B-conjugated PE or PS liposomes were incubated with ATG4B/C/D or RavZ for 60 mins and analysed by SDS-PAGE/Coomassie and **f**, Densitometry. Data from 2 independent experiments normalized to untreated controls. **g**, GFP-LC3B was conjugated to PE and PS on mixed liposomes, incubated with ATG4s or RavZ for 60 mins then analysed by mass spectrometry. Data represent means normalised to untreated controls from 3 independent experiments, ***p<0.0004, **p<0.004, *p<0.01, paired t test. **h-i**, Wild type and *ATG4D*^*-/-*^ HCT116 cells, under **h**, basal or **i**, monensin treated conditions, analysed for GFP-LC3B lipidation by mass spectrometry. Data are means normalised to wild type levels from 2 independent experiments.

However, SMAC is a transient and reversible process ^24^, inferring that delipidation of both ATG8-PE and ATG8-PS components is likely to occur in cells. We therefore reasoned that an alternative ATG4 isoform may catalyse PS delipidation. To test this, a panel of ATG4 proteins (B, C, D) were purified, from mammalian cells, and their delipidation profiles determined in liposome-based assays (Fig. 4e, f). We note that full length ATG4C and ATG4D are active under these conditions, without caspase cleavage, unlike the bacterially purified proteins ^36,37^, and ATG4A was omitted, as it appears GABARAP specific ^36,38^; RavZ, a bacterial effector protein known to cleave both ATG8-PE and PS ^39,40^, was included as a control. As expected, all 4 enzymes can delipidate LC3B from PE liposomes, consistent with previous work ^36^. Strikingly however, only ATG4D (and the RavZ control) can deconjugate LC3B from PS liposomes. To control for any indirect effects of differing liposome composition, a mixed lipid system was also assessed, where LC3B is conjugated to PE or PS on the same liposomes. Here too, LC3B-PS is more efficiently delipidated by ATG4D than ATG4B, while LC3-PE is deconjugated by both (Fig. 4g). These data indicate that ATG4D may represent the major ATG8-PS delipidating activity. Consistent with this, CRISPR deletion of *ATG4D* elevates cellular levels of both LC3-PS and LC3-PE, under both basal (Fig. 4h) and monensin-induced conditions (Fig. 4i).

Together, these findings suggest that ATG4D plays a major role in delipidating ATG8-PS in the non-canonical autophagy/SMAC pathway, providing new insights into ATG4 function and isoform specificity. Notably, while ATG4 proteolytic activity is quite promiscuous, with many amino acids accommodated downstream of the scissile Gly ^35^, delipidating activity appears much more selective due to the structural constraints of the lipid headgroup.

The C-terminal lipidation of ATG8 is a unique post-translational modification and a hallmark event during autophagy related processes, widely used to detect and monitor the pathway. Here, we report the first cellular evidence for alternative ATG8 lipidation to PS. Alternative ATG8 lipidation occurs upon activation of non-canonical autophagy, during pharmacological SMAC, LC3-associated phagocytosis and Influenza A infection, at single-membrane endolysosomal compartments, but is not observed during canonical autophagy. Unlike ATG8-PE, ATG8-PS is selectively deconjugated by ATG4D, revealing important molecular distinctions between the different forms of ATG8-II and new insights into ATG4 family functions. Finally, ATG8-PS provides a ‘molecular signature’ for the non-canonical autophagy/SMAC pathway and opens up a range of new mechanistic and functional questions for future study.

---

**Extended Data Figure 1:**
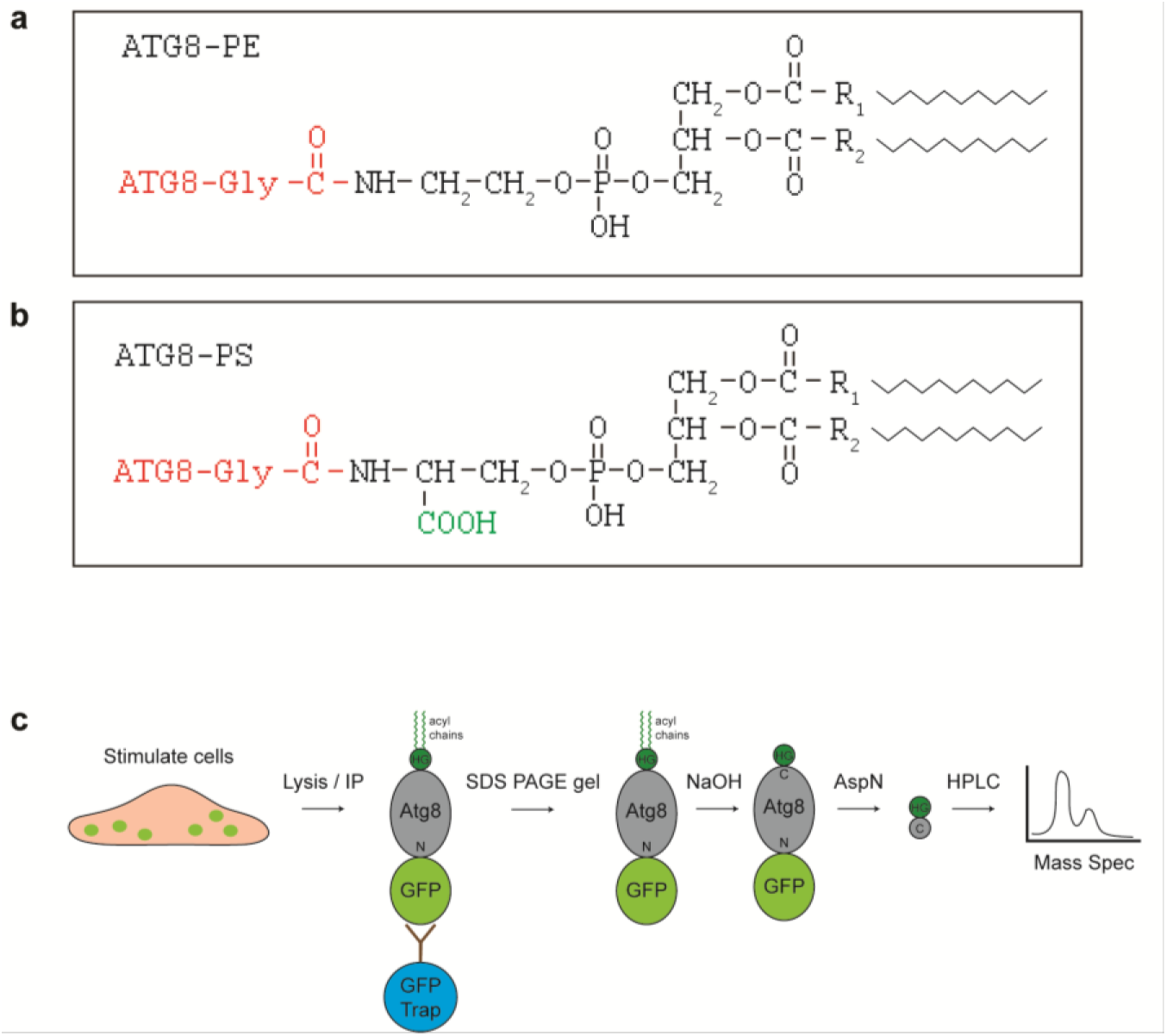
Analysis of ATG8 lipidation. Schematics representing ATG8 conjugated to **a**, PE and **b**, PS headgroup moieties. **c**, Work flow for GFP-ATG8 mass spectrometry analysis. Cells are stimulated as indicated, then subjected to lysis and GFP-ATG8 immunoprecipitation using GFP-TRAP beads. Samples are run on SDS-PAGE gels, coomassie stained and appropriate bands excised and base treated to strip lipid acyl chains. HG = lipid headgroup. Samples are then digested using AspN protease and analysed by mass spectrometry.

**Extended Data Figure 2:**
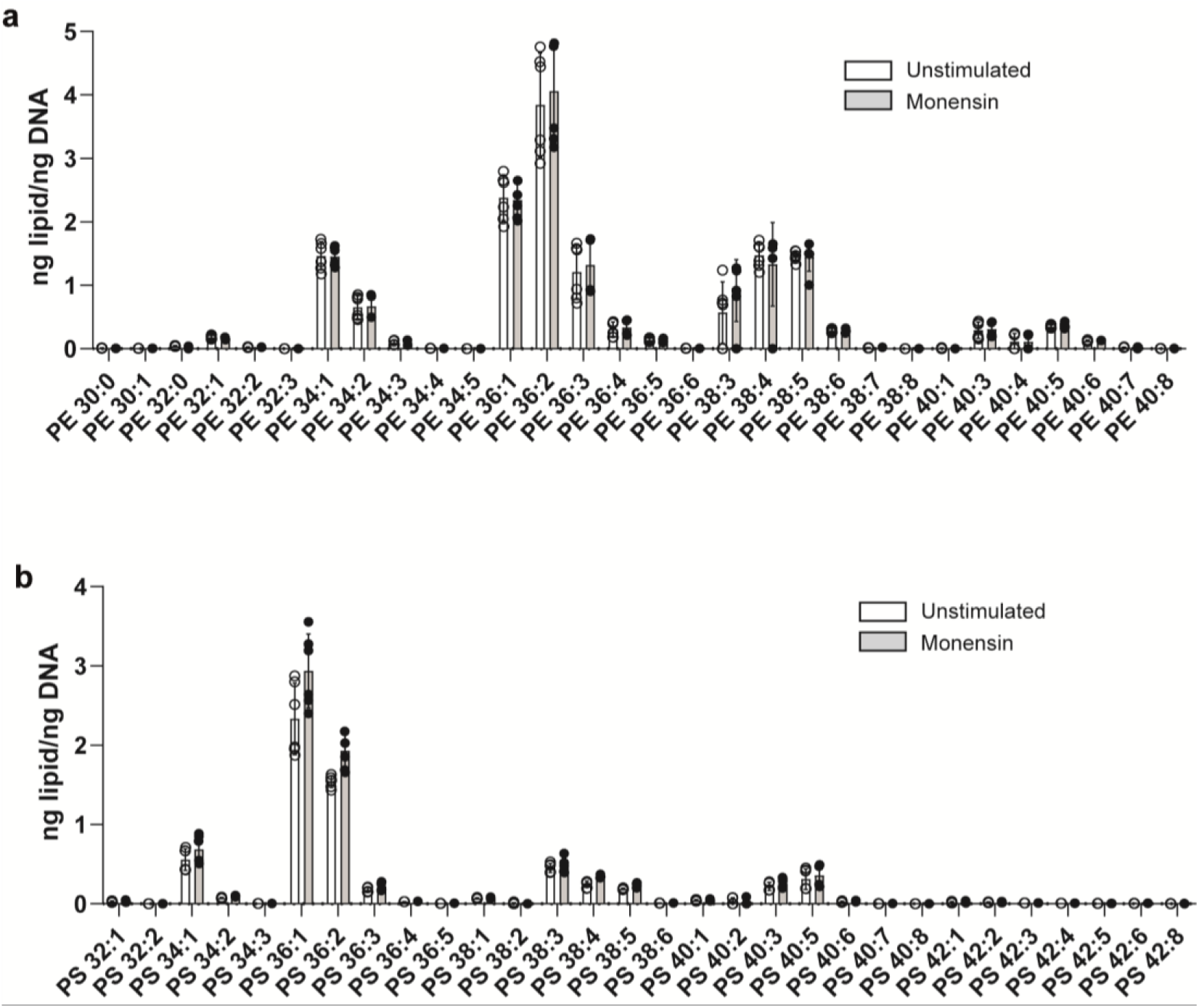
Lipidomic analysis of monensin treated cells. Global lipids from *ATG13*^*-/-*^ MCF10A cells +/-100 μM monensin for 60 mins were analysed for **a**, phosphoethanolamine (PE) or **b**, phosphatidylserine (PS) molecular species. Data represent means from 6 biological replicates.

**Extended Data Figure 3:**
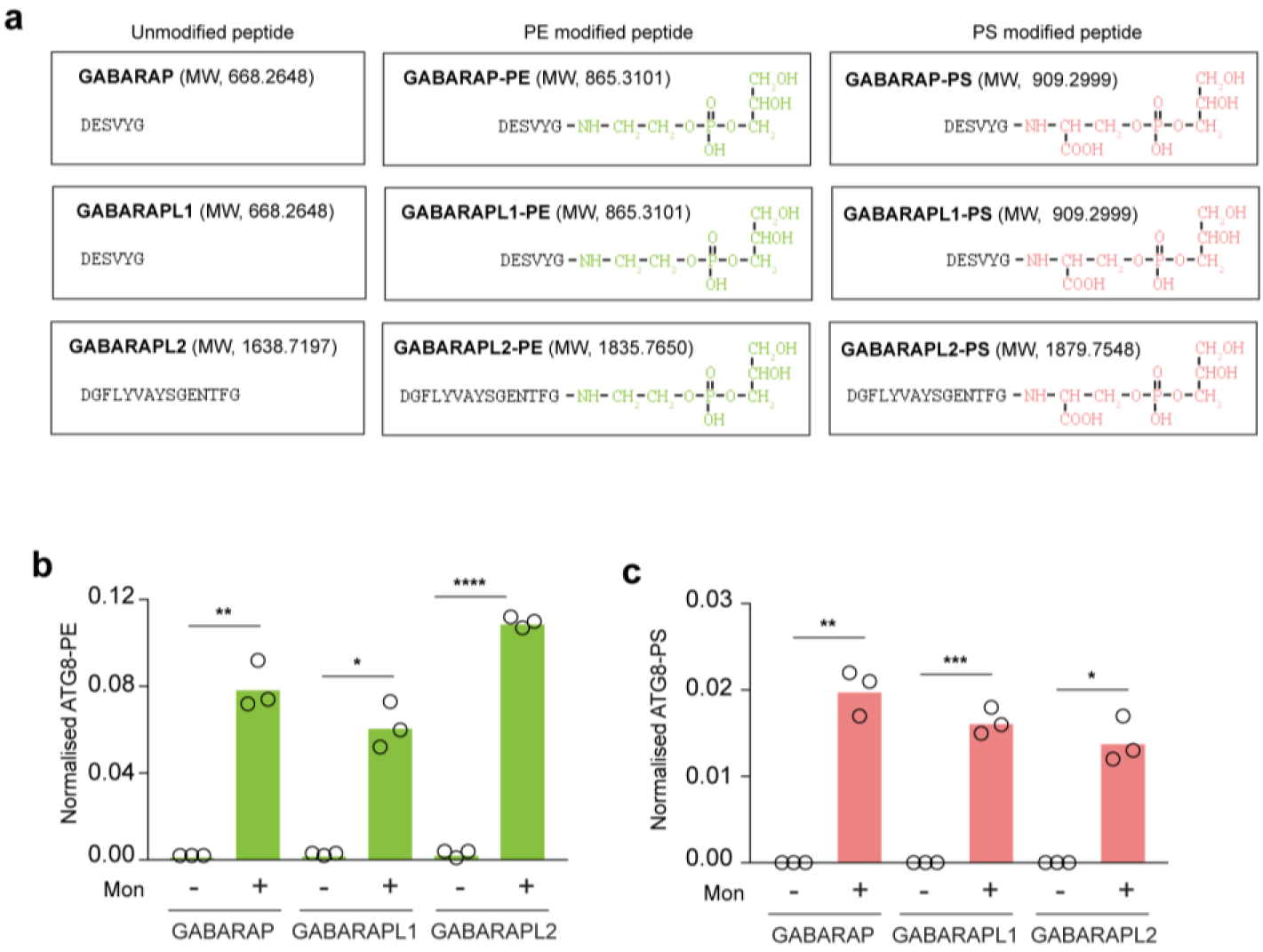
GABARAP proteins are conjugated to PS during non-canonical autophagy. **a**, C-terminal peptides of GABARAP proteins conjugated to either the PE or PS headgroup. Predicted MWs are indicated. **b, c**, MCF10A cells expressing different GFP-tagged GABARAP proteins were treated with 100 μM monensin for 60 mins to activate SMAC and analysed for lipidation to PE and PS. Data represent means from 3 independent experiments, ****p<0.0003, ***p<0.003, **p<0.007, *p<0.01, paired t test.

**Extended Data Figure 4:**
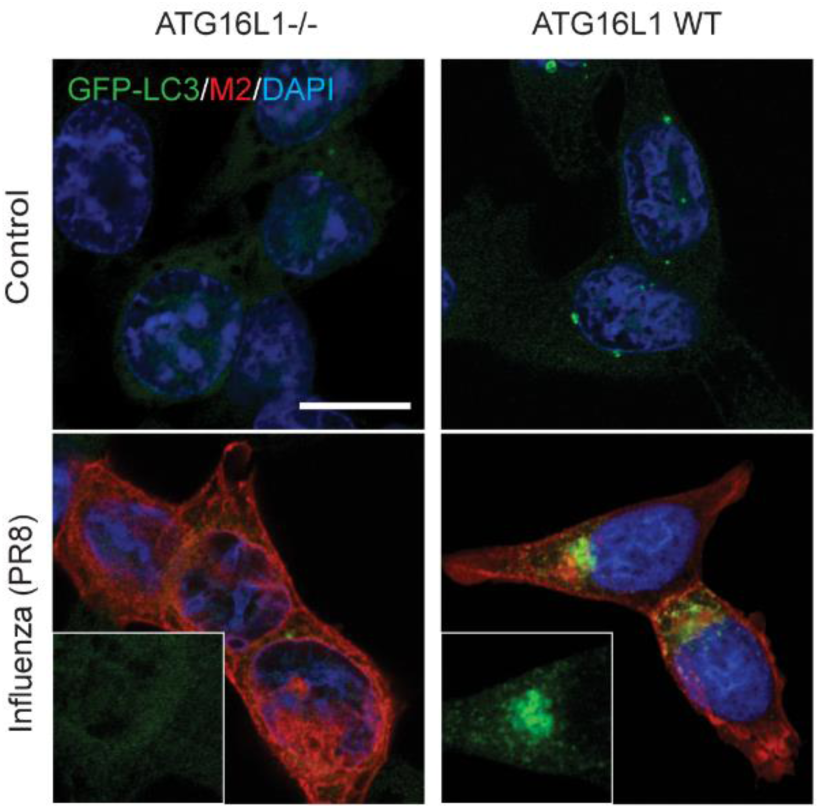
Influenza A infection induces GFP-LC3B relocalisation. Representative confocal images of HCT116 *ATG16L1*^*-/-*^ cells and ATG16L1 WT re-expressing cells infected with PR8 (m.o.i. 5) for 16 hours. Merged images of GFP-LC3 (green), viral M2 (red) and DAPI (blue). Cropped images of GFP-LC3 in PR8 stimulated cells. Scale bar: 10 μm.

**Extended Data Figure 5:**
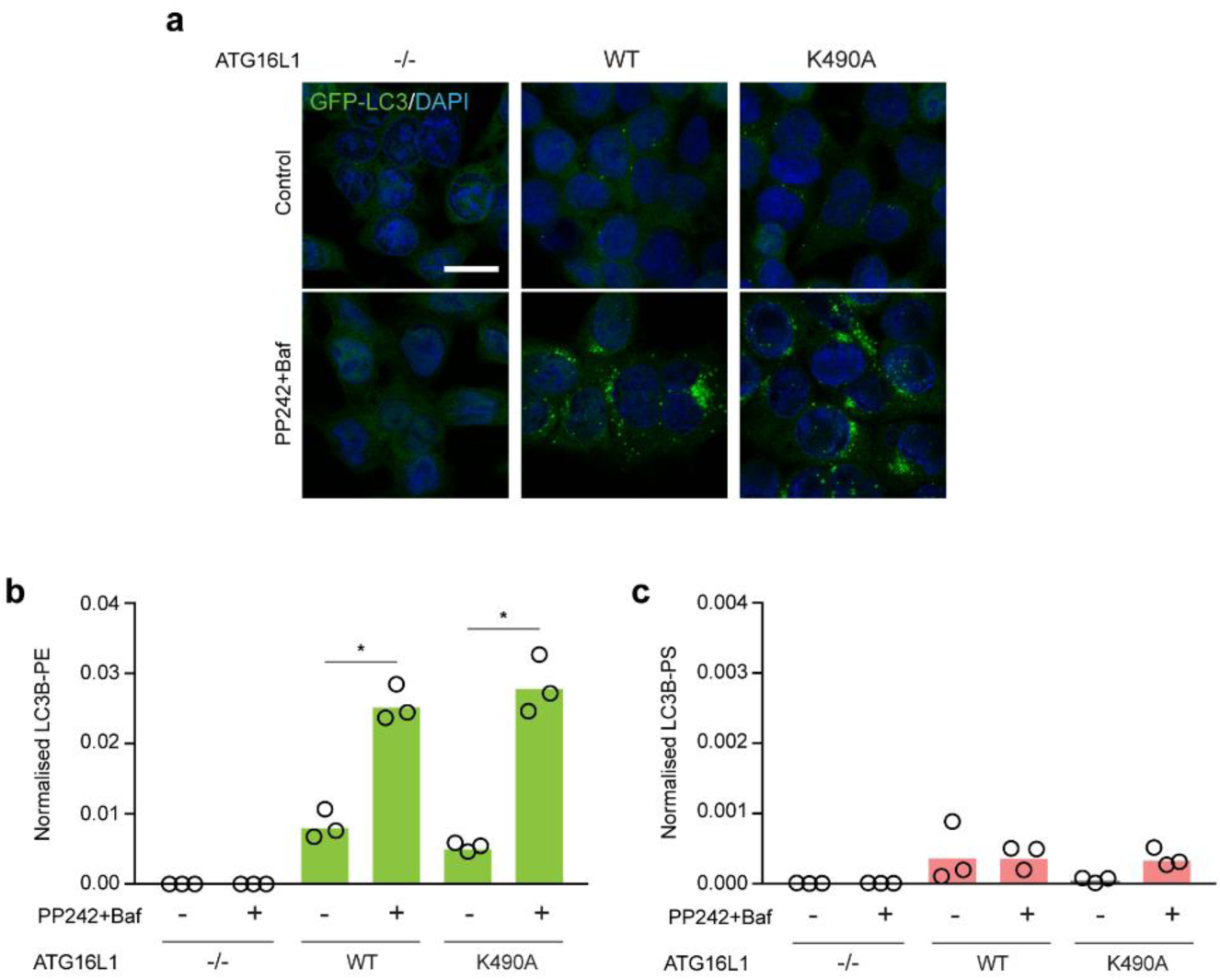
ATG16L1 K490A supports efficient LC3-PE lipidation during canonical autophagy. HCT116 *ATG16L1*^*-/-*^ cells, re-expressing ATG16L1 WT or K490A, and GFP-LC3B, were stimulated -/+ 1 μM PP242/100 nM BafA1 for 60 mins. **a**, Merged confocal images of GFP-LC3B (green) and DAPI (blue). Scale bar: 10 μm. **b, c**, Mass spectrometry analysis of LC3B conjugation to PE and PS. Data represent means from 3 independent experiments, *p<0.01, paired t test.

**Extended Data Figure 6:**
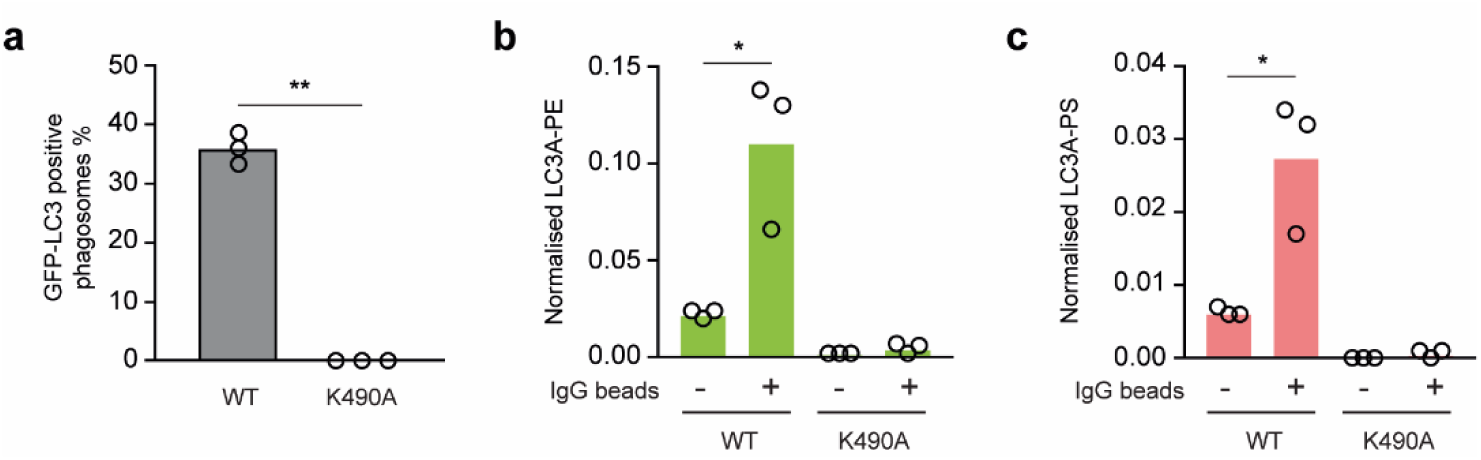
LC3A-PS lipidation is dependent on the ATG16L1 WD40 domain during LAP. **a**, Quantification of GFP-LC3A recruitment to IgG-bead containing phagosomes in RAW264.7 ATG16L1-/-cells re-expressing WT or K490A ATG16L1. Data represent means from 3 independent experiments, WT n=189, K490A n=150 phagosomes in total, **p<0.002, paired t test. **b, c**, Mass spectrometry analysis of LC3A conjugated to PE or PS during LAP in RAW264.7 cells expressing ATG16L1 WT or K490A. Data represent means from 3 independent experiments, *p<0.01, ratio paired t test.

**Extended Data Figure 7:**
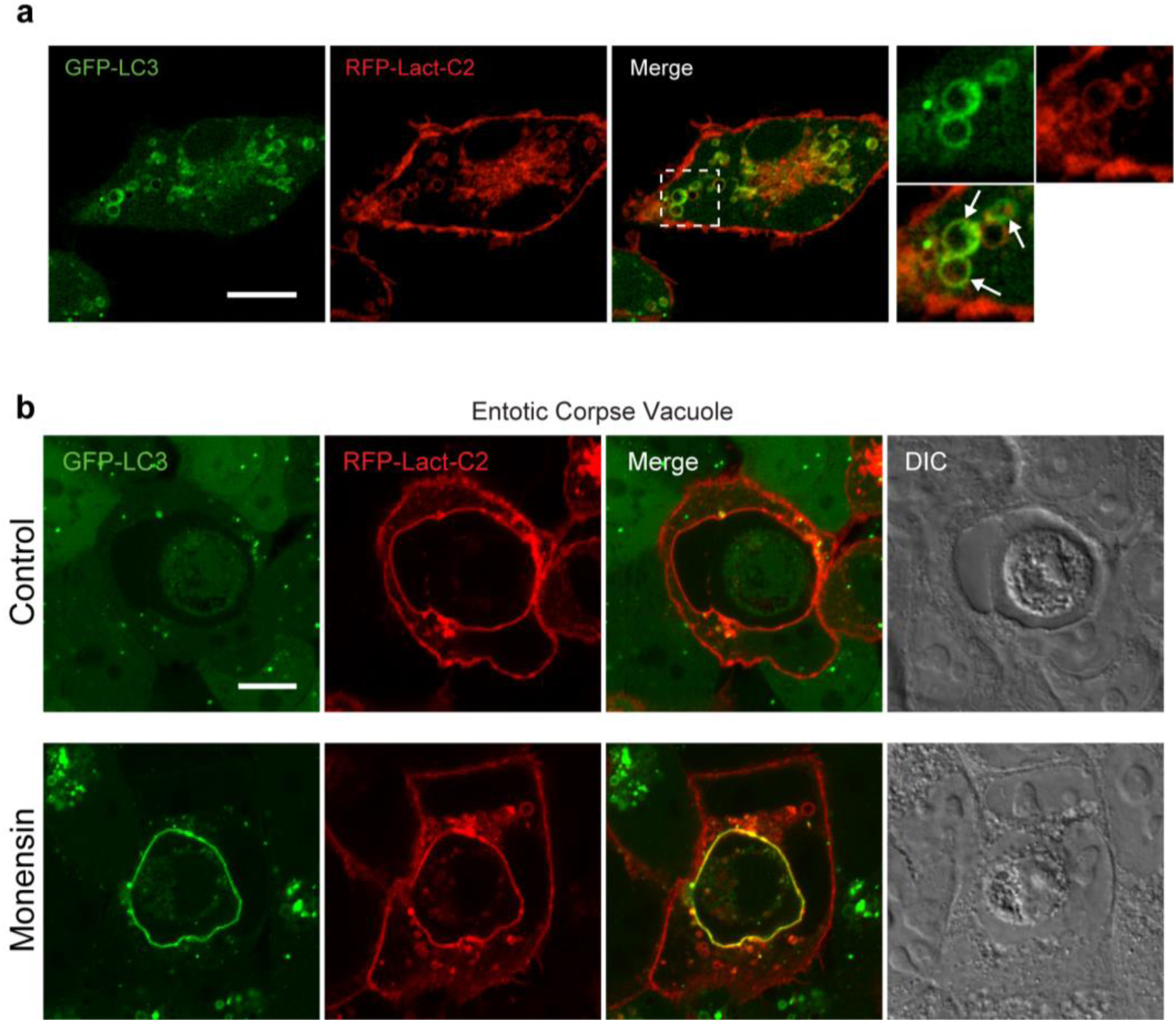
GFP-LC3A is recruited to PS positive macropinosomes and entotic corpse vacuoles. **a**, Representative confocal images of J774A.1 cells expressing GFP-LC3A and PS sensor RFP-Lact-C2. Arrows denote macropinosomes. Scale bar: 5 μm. **b**, Confocal images of entotic corpse vacuoles in MCF10A expressing GFP-LC3A and RFP-Lact-C2. Scale bar: 10 μm.

## Methods

### Reagents

Bafilomycin A1 (#1334) and PP242 (#4257) were purchased from Tocris; Monensin (M5273), DAPI (D9542) and human IgG (I4506) were from Sigma. GFP-Trap (gtma-20) and control magnetic agarose beads (bmab-20) were obtained from Chromotek, Magnetic 3-micron beads (PMA3N) from Bangs Laboratories and Latex polymer 3-micron beads (17134-15) from Polysciences. Murine IFNγ (315-05) was from Peprotech. Lipids were purchased from Avanti Polar Lipids (Alabaster, AL), dissolved in chloroform: 1,2-dioleoyl-sn-glycero-3-phosphoethanolamine (DOPE; 850725C), 1-palmitoyl-2-oleoyl-sn-glycero-3-phosphocholine (POPC; 850457C), 1,2-dioleoyl-sn-glycero-3-phospho-L-serine (DOPS; 840035C) and 1,2-dioleoyl-sn-glycero-3-phosphoethanolamine-N-(lissamine rhodamine B sulfonyl) (Liss Rhod PE; 810150C).

### Plasmids

GFP-tagged, human GABARAP, GABARAPL1 and GABARAPL2, in pBabe-Puro, were purchased from MRC-PPU, University of Dundee. mRFP-Lact-C2 was a gift from Sergio Grinstein (Addgene plasmid # 74061). GFP-huLC3A pBabe-Blast was kindly provided by Dr Michael Overholtzer (MSKCC). Flag-S-tagged versions of mouse ATG16L1 (wild type and K490A mutant), in pBabe-Puro, were previously described ^8^.

### Cell Lines and Culture

WT or *ATG13*^*-/-*^ MCF10A cells (human breast epithelial), expressing GFP-LC3A (human), were prepared as described previously ^5^ and cultured in DMEM/F12 (Gibco, 11320074) containing 5% horse serum (Sigma), EGF (20ng/ml; Peprotech AF-100-15), hydrocortisone (0.5 mg/ml; Sigma, H0888), cholera toxin (100 ng/ml; Sigma, C8052), insulin (10 μg/ml; Sigma, I9278), and penicillin/streptomycin (100 U/ml, /ml; Gibco 15140-122) at 37°C, 5% CO_2_. These parental cell lines were also engineered to express alternative GFP-tagged isoforms of human ATG8s, using retroviral infection (pBabe-Puro) and antibiotic selection (2.5 μg/ml Puromycin). HCT116 cells (human colorectal epithelial) were maintained using DMEM (Gibco, 41966-029) supplemented with 10% FBS (Sigma) and penicillin/streptomycin (100 U/ml, 100 μg/ml; Gibco 15140-122) at 37°C, 5% CO_2_. A panel of lines expressing GFP-LC3B (rat) and different ATG16L1 constructs, were derived from *ATG16L1*^*-/-*^ cells, reconstituted with the pBabe-Puro *ATG16L1* (wild type or K490A), as described previously ^8^. ATG4D null cells were prepared as described below.

J774.A1 mouse macrophage were obtained from ATCC and cultured in DMEM (Gibco, 41966-029) supplemented with 10% FBS (Sigma) and penicillin/streptomycin (100 U/ml, 100 μg/ml; Gibco 15140-122) at 37°C, 5% CO_2_. These cells were engineered to express GFP-LC3A (human) by retroviral infection (pBabe-Blast) and antibiotic selection (8ug/ml Blasticidin).

*ATG16L1*^*-/-*^ RAW264.7 mouse macrophage were generated as previously described ^26^ and cultured in DMEM (Gibco, 41966-029) supplemented with 10% FBS (Sigma) and penicillin/streptomycin (100 U/ml, 100 μg/ml; Gibco 15140-122) at 37°C, 5% CO_2_. These cells were engineered to express GFP-LC3A (human, pBabe-Blast), and reconstituted with *ATG16L1* wild type or K490A (pBabe-Puro), all by retroviral infection and selection (8 μg/ml Blasticidin, 2 μg/ml Puromycin). HEK293FT cells were grown in DMEM (Gibco, 41966-029) supplemented with 10% FBS (Sigma) and penicillin/streptomycin (100 U/ml, 100 μg/ml; Gibco 15140-122) at 37°C, 5% CO_2_.

### Generation of ATG4D CRISPR knock out cells

Stable knock out cell lines were generated using CRISPR technology by nucleofection of HCT116.GFP-LC3B cells with a pool of *in vitro* synthesised guide RNAs (Synthego) and Cas9 (Thermo). Single cell clones were isolated and absence of gene expression confirmed by western blotting. The sgRNAs were designed using the Synthego software: ATG4D guide 1: ggcgggacacaaagucccgc, ATG4D guide 2: gggacuuugugucccgccug, ATG4D guide 3: cccggcgguaugugagccac.

### Retrovirus Production and Infection

Retrovirus production and infection was performed as described previously in HEK293T cells ^41^. Selection was achieved with antibiotic treatment for 2-5 days. Constructs, plasmids and antibiotic concentrations are all indicated above.

### Pharmacological Stimulation

To induce canonical autophagy, cells were pretreated for 20 mins with 100 nM bafilomycin, followed by addition of 1 μM PP242 for a further 40mins. To induce non-canonical autophagy/SMAC, cells were treated with 100 μM monensin for 60 mins. Stimulated cells were analysed by microscopy, or lysed for western blotting or mass spectrometric analysis, as indicated.

### J774.A1 Phagosome Preparation and Assay

To induce, enrich and analyse phagosomes, J774.A1 cells expressing GFP-LC3A (human) were assayed with IgG coated magnetic beads (ProMag 3 Series-Amine, Bangs Laboratories). The magnetic beads were prepared according to the manufacture’s guidelines. Briefly, beads were: i) washed in PBS and activated by rotating with 10% gluteraldehyde for 1 hour, RT; ii) washed in PBS and resuspended by rotating with 6 mg human IgG (Sigma, I4506) for 2 hours, RT; iii) washed again and quenched by rotating with 40 mM glycine for 1 hour, RT and iv) finally resuspended in PBS.

To enrich phagosomes for LC3 lipidation analysis, 8 × 15cm plates of J774.A1 cells were seeded per condition, incubated for 3 days, then stimulated with 200 U/ml murine IFNγ (Peprotech, 315-05) for 24 hours. Cells were then preincubated with 100 nM Bafilomycin A1, or DMSO control, for 15 mins, then phagocytosis was induced with IgG coated beads, which were incubated for 25 mins, 37°C. Cells were placed on ice and washed with ice cold PBS. Each dish was scraped into 0.5 ml HB buffer: 250 mM sucrose, 10 mM HEPES, phosphatase inhibitors (1x, Sigma P0044) and protease inhibitors (1x, Sigma P8340), then spun at 200 rcm, 5 mins. The pellet (containing intact cells and beads) was resuspended in 1 ml fresh HB buffer and an aliquot of total cell extract removed. Cells were then gently ruptured with 35 strokes of a Dounce homogenizer, on ice. Samples were placed on a magnetic rack, to isolate the magnetic beads and their enclosing phagosomes. The beads were washed with 2x 1 ml HB buffer and the parallel samples for each condition pooled; an aliquot of this phagosome preparation was withheld. Finally, to release and recover the phagosomal GFP-LC3 for analysis, the bead pellet was lysed in NP40 lysis buffer and subjected to GFP-TRAP IP, as described below.

### RAW264.7 Phagocytosis Assay

IgG-coated latex beads were prepared as previously described ^42^. Briefly, 3-micron beads (Polysciences Inc) were resuspended in 0.1 M Borate and incubated with human IgG at 4°C overnight while rotating. Beads were washed in PBS x 3 before resuspension in PBS. RAW2647 cells were seeded in 15 cm^2^ dishes and treated with 200 U/ml IFNγ (Peprotech, 315-05) for 24 hours prior to use. Where indicated, 350 ul IgG beads were added to dishes for 30 minutes at 37°C. Cells were washed in cold PBS x 1 and lysed in 900 ul lysis buffer consisting of: 50 mM Tris pH 7.5, 150 mM NaCl, 0.5% NP40 (IGEPAL CA-630, Sigma I3021), phosphatase inhibitors (1x, Sigma P0044) and protease inhibitors (1x, Sigma P8340). Samples were scraped into pre-chilled 1.5 ml Eppendorf tubes. Samples were incubated on ice for 20 minutes and centrifuged at 13,500 rpm for 10 minutes at 4°C. The supernatants were kept and subjected to GFP-TRAP IP, as described below.

### Influenza A Infection

Stocks of influenza A virus PR8 (strain A/Puerto Rico/8/1934) were generated using an eight plasmid-based system, as previously described ^43^, and propagated on MDCK cells. For infection, cells were washed with serum-free DMEM, then incubated with virus in serum-free DMEM at 37°C. After 1 h, the medium was replaced with DMEM containing 10% FBS. Cells were processed 16 hours post infection (h.p.i.) and analysed by microscopy, or lysed for western blotting or mass spectrometric analysis, as indicated.

### Whole Cell Lipidomic Analysis

5 × 10^5^ MCF10A cells were seeded per 6 cm dish, incubated overnight and then treated as indicated. These cells adhere strongly and tend to rupture upon scraping, so were harvested by trypsinsation. Cells were rinsed in PBS, incubated with trypsin for 3 minutes, 37°C and harvested in media. The cells were then washed 3x in PBS and pelleted at 150 rcf, 3 minutes. Cell pellets were snap frozen in liquid nitrogen for lipid analysis. The frozen cell pellets were subjected to Folch extraction using chloroform/MeOH/H_2_O (2:1:1). The dry extract was re-suspended in chloroform/MeOH (1:1). Phosphatidylethanolamine (PE) and phosphatidylserine (PS) were separated by liquid chromatography using a Shimadzu XR system (Shimadzu, Kyoto, Japan) ^44^. PE and PS were then detected using an Orbitrap Elite mass spectrometer in full scan mode with a mass range of 400-1000 *m/z* at a target resolution of 240,000 (FWHM at *m/z* 400). Data were analysed using Lipid Data Analyzer (2.6.0–2) software ^45^.

### Cell Lysis and GFP-TRAP Immunoprecipitation

Cells expressing GFP-ATG8 were seeded across multiple 15-cm dishes, treated as indicated, then placed on ice and washed with ice-cold PBS. Each 15-cm dish was scraped into 900 μl lysis buffer. Lysis composition was as follows: 1) GFP-LC3A/B in MCF10A/RAW264.7: 50 mM Tris pH 7.5, 150 mM NaCl, 0.5% NP40 (IGEPAL CA-630, Sigma I3021), phosphatase inhibitors (1x, Sigma P0044) and protease inhibitors (1x, Sigma P8340); 2) GFP-LC3B in HCT116 cells, as above, except 1% Triton replaces 0.5% NP40; 3) GFP-GABARAPs in MCF10A: as above, but with the addition of 20 mM N-Ethylmaleimide (NEM) to protect the lipidated species ^46^. The resulting suspension was incubated on ice for 10 minutes, then centrifuged at 16000 rcm, 4°C, 10 minutes to separate the pellet from the soluble lysate. A small fraction of the supernatant was removed for western blotting, as described below, and the remaining lysate subjected to preclearing and IP, using magnetic beads (Chromotek) and a magnetic separation rack (Cell Signalling), according to the manufacturers’ instructions. The lysate was pre-cleared, using 10 μl equilibrated magnetic agarose control beads/sample (bmab, Chromotek), for 30 minutes, 4°C, on a rotating wheel. Cleared lysates were then incubated with 10 μl equilibrated GFP-TRAP beads/sample (gtma, Chromotek) for 60 minutes, 4°C, on a rotating wheel, to recover GFP-LC3. The beads were washed 3 × 10 minutes in lysis buffer at 4°C, on a rotating wheel. Enriched GFP-LC3 was either processed further on the beads (see ATG4B delipidation assay), or eluted for analysis by Mass Spectrometry with the addition of 25 μl 2x LDS buffer (Invitrogen)/0.2 M DTT sample buffer at 100°C, 5 minutes.

### Mass Spectrometric Analysis of Lipidated ATG8

ATG8 samples were run on 10% NuPAGE gels in MOPS buffer (Invitrogen), alongside protein molecular weight markers (EZ-Run, Fisher). Gels were released into a MeOH rinsed box for washing and staining, all at RT, with gentle shaking. Each gel was washed 3 × 5 mins in dH2O, stained with Imperial Stain (Thermo Scientific, 24615) for 2 hrs, then destained in dH2O overnight. Stained gels were scanned and representative images are presented. For each sample, the entire gel region containing both lipidated and non-lipidated ATG8 protein was excised into a single tube, destained, and typically saponified by treatment with 50 mM NaOH in 30% MeOH at 40°C for 2 hr. The protein was digested with AspN protease (Roche) at 30°C for 16 hr, in 25 mM ammonium bicarbonate, which cleaves predominantly to the N-terminal side of Asp residues.

For the initial characterisation of modified LC3A, peptides were separated on a reversed-phase nanoLC column (150 × 0.075mm; Reprosil-Pur C18AQ, Dr Maisch), interfaced to an Orbitrap Velos Pro mass spectrometer (Thermo Scientific), operating in high resolution (orbitrap) MS1 mode, with data-dependent acquisition of low resolution MS2 spectra generated by CID in the linear ion-trap. The measured neutral monoisotopic masses of the three forms of LC3A C-terminal peptide DGFLYMVYASQETFG, calculated from the predominant doubly protonated pseudomolecular ions, were: unmodified - 1726.758 (theoretical 1726.754); PE-modified - 1923.802 (theoretical 1923.800); PS-modified - 1967.790 (theoretical 1967.789). MS2 data were searched against the Uniprot mouse proteome database using Mascot software (Matrix Science), with glycerophosphoethanolamine and glycerophosphoserine combined with loss of the C-terminal amino-acid, specified as custom C-terminal variable modifications. Spectra which were matched to the C-terminal modified peptides were confirmed by manual interpretation.

For the targeted mass spectrometric assay of modified C-terminal LC3A/B peptides, samples were processed identically, but the analysis was done on a Q-Exactive mass spectrometer (Thermo Scientific). It was observed during the characterization of the C-terminal peptides that the Met residue was >90% oxidized to the sulphone, so in order to increase the sensitivity of the analyses, the Met-oxidised forms of the three peptides were targeted. The mass spectrometer scan cycle consisted of one high-resolution MS1 scan, and three high resolution MS2 scans from fragmentation of the doubly charged parent ions of the unmodified, glycerophosphoethanolamine- and glycerophosphoserine-modified ATG8 C-terminal peptide. Quantitative data were extracted using Skyline software (MacCoss Lab, University of Washington) using the sum of the chromatographic peak areas from the y1 to y10 fragment ions. Pilot studies showed that modified peptide peak area could be normalised against unmodified C-terminal peptide, a distinct internal LC3 peptide or a GFP-tag derived peptide, with similar results. Subsequently, normalization was performed against unmodified C-terminal peptide, unless otherwise indicated.

The GABARAP proteins were analysed only in targeted mode, analogously to LC3A/B. The C-terminal peptide for the GABARAPL2, DGFLYVAYSGENTFG, is similar to the LC3A/B peptide and is predominantly doubly charged. The GABARAP and GABARAPL1, C-terminal peptide DESVYG is predominantly singly charged.

### On bead ATG4B Delipidation Assay

*ATG13*^*-/-*^ MCF10A cells were treated -/+ 100 μM monensin for 60 minutes and GFP-LC3A was enriched, immobilised and washed on GFP-TRAP beads, as described above. Recombinant His-tagged human ATG4B (Abcam, ab188707) was pre-treated with 10 mM DTT for 15 mins, RT, to achieve maximum activation, then added at 2 μg/sample, in lysis buffer, to the GFP-LC3 beads for 0-120 minutes at 37°C. At the end of the time-course, the reaction mixture was aspirated, the beads quickly rinsed with ice cold lysis buffer and GFP-LC3 was eluted for analysis by Mass Spectrometry with 25 μl 2x LDS (Invitrogen)/0.2 M DTT sample buffer at 100°C, 5 mins.

### Protein Purification

Human ATG3, ATG7, and ATG12–ATG5-ATG16L1ß complex, were expressed and purified from HEK suspension cells as described ^26^. Human ATG4B, ATG4C, and ATG4D were expressed in HEK suspension cells from pCMV plasmids encoding full-length proteins using the following constructs: GST-[HRV 3C protease cleavage site]-3xFlag-[TEV protease cleavage site]-ATG4B, 3xFlag-[SUMOstar protease cleavage site]-ATG4C, GST-[HRV 3C protease cleavage site]-ATG4D-[TEV protease cleavage site]-3xFlag. Cells at 2 × 10^6^ cells/ml in BalanCD medium (Irvine Scientific) were transfected with 1 microgram plasmid/10^6^ cells in 100-ml batches, using polyethyleneimine as transfection agent (3 microgram /10^6^ cells), and incubated by horizontal shaking (160 rpm) at 37°C in 8% CO_2_ for 2 days with addition of 5% Feed (Irvine Scientific) after 1 day. Cells were harvested by centrifugation and lysed in 1% Nonidet P40/1 mM EDTA/PBS containing Complete protease inhibitors (Roche), and nuclei were removed by centrifugation. After an additional centrifugation at 20.000 x g for 10 minutes, ATG4B in the supernatant was bound to glutathione-Sepharose (GE Healthcare), washed with NT350, and eluted by on-column cleavage with TEV protease at 4°C overnight. Eluted protein was gel filtrated on Sephacryl S-300 HR (GE Healthcare) equilibrated with NT350, 1 mM DTT. Eluted fractions with purified ATG4B were pooled, snap-frozen and stored at −80°C. ATG4C and ATG4D were purified by the same procedure, but with the following differences: ATG4C was bound to M2-agarose (Sigma) and eluted by on-column cleavage with SUMOstar protease. ATG4D was bound to M2-agarose, and after elution with TEV protease the protein was bound to glutathione-Sepharose and eluted with HRV 3C protease.

LC3B (amino acids 1-120) was expressed in BL21 pLysS DE3 *E*.*coli* cells from a pGEX-6P-2 plasmid (GE Healthcare) in 250 ml LB medium. After induction with IPTG at OD 0.8, the cells were grown at 22°C for 4 hours and harvested by centrifugation. After washing with NH100 (100 mM NaCl, 20 mM HEPES-KOH pH 7.4), the cells were resuspended in 5 ml NH100 and snap-frozen. The cells were thawed and centrifuged at 75.000 x g for 30 minutes, and the supernatant was incubated with glutathione-Sepharose. After washing with NH100, LC3B was eluted by on-column cleavage overnight at 4°C with HRV 3C protease. Eluted protein was gel filtrated on Sephacryl S-200 HR (GE Healthcare) equilibrated with NH100 buffer, and purified LC3B was snap-frozen and stored at −80°C.

RavZ was expressed in BL21-Gold (DE3) *E. coli*. Cells were grown at 37°C to an OD of 0.6–0.8 before protein expression was induced with 0.5 mM IPTG. Cells were then grown for three additional hours before they were collected by centrifugation. Cells were resuspended in NT350 supplemented with a Roche Complete Protease Inhibitor, lysed by sonication, and cleared by centrifugation (20.000 x g for 10 minutes). The supernatant was incubated at 4°C with Glutathione Beads (Sigma) for 4 hours. Beads were collected and washed twice with NT350 buffer before HRV 3C protease was added and allowed to cut at 4°C overnight. The next morning, protein fractions were collected and stored at −80°C.

### Liposome Assays

To prepare liposomes the various lipid combinations were dried under nitrogen gas, and the lipid film was further dried under vacuum for 1 hour. The lipids were reconstituted in NT350 buffer (350 mM NaCl, 20 mM Tris–HCl pH 7.4) and subjected to seven cycles of flash-freezing in liquid nitrogen and thawing in a 37°C bath. Liposomes were further sonicated immediately prior to the lipidation reaction. Lipidation reactions were carried out in microcentrifuge tubes containing ATG7 (0.5 µM), ATG3 (1 µM), ATG12-ATG5-ATG16L1 (0.05 µM), LC3B (aa1-120, 10 µM) and sonicated liposomes (3 mM), mixed in NT350 buffer containing 1 mM DTT. Lipidation was initiated by adding 1 mM ATP and reactions were incubated at 37°C for 90 minutes. The lipidation reaction was then run on a Nycodenz density gradient to remove non-lipidated LC3B and other reaction components from the proteoliposomes. The bottom layer of the gradient consisted of 150 µL of 80% Nycodenz and 150 µL of the lipidation reaction. The second layer consisted of 250 µL of 30% Nycodenz while the top layer was 50 µL of NT350 buffer. Gradients were spun at 48000 rpm at 4°C for 4 hours in a Beckman SW55Ti rotor. Liposomes with the lipidated LC3B was collected from the top of the tube and stored at 4°C before use in subsequent de-lipidation experiments.

To measure de-lipidation of LC3B-PE/LC3B-PS, proteoliposomes (∼1 µM LC3B-II) were mixed with NT350 containing 1 mM DTT and kept on ice until activity assays were initiated by addition of various ATG4 proteins (0.1 µM) or RavZ (0.1 µM).

Reactions were incubated at 30°C for 1 hour, before they were mixed with sample buffer and immediately boiled to stop proteolysis. The samples were then separated using SDS-PAGE, visualised by Coomassie staining and either analyzed directly with ImageLab (Biorad) to assess bandshift, or processed for Mass Spectrometry, as described above. To quantify levels of delipidation, densitometry LC3I and LC3II in Coomassie images was performed using Image J.

### Western Blotting

Western blotting was performed as described previously ^5,8^. Briefly, cell lysates were run on SDS-PAGE gels (10%, 12% or 15%), transferred to PVDF membrane (Immobilon-P, Millipore), blocked with 5% BSA (Sigma A7906)/TBS-T for 1 hour, RT and then incubated with primary antibody at 4°C overnight. The following antibodies were used, all at 1:1000: anti-ATG13 (Cell Signaling, 13272), anti-ATG16L1 (Cell Signaling, 8089), anti-GAPDH (Abcam, ab9843), anti-GFP (Sigma, 11814460001) and anti-M2 (Abcam, ab5416). Membranes were washed 3x 10 minutes in TBS-T and incubated with HRP-conjugated secondary antibodies (Cell Signalling 7074, 7076) for 45 minutes, RT. Membranes were washed again 3x 10 minutes in TBS-T, then developed with ECL (GE, RPN2209). Blots were scanned (Epson Perfection, V550) and images are representative of 3 separate experiments.

### Microscopy

For fixed cell imaging cells were washed in PBS x 1 followed by fixation in cold methanol at −20°C for 5 minutes. Samples were washed with PBS and blocked in 5% BSA/PBS for 1 hour at room temperature. Where indicated, primary antibodies were diluted in blocking buffer and added overnight at 4°C. Cells were washed and secondary antibody diluted in blocking buffer added for 1 hour. Final washes were performed, before incubating with DAPI for 10 minutes and mounting in ProLong Gold Antifade (ThermoFisher, P36930). Samples were analysed on a Confocal Zeiss LSM 780 microscope (Carl Zeiss Ltd) equipped with a 40x oil immersion 1.40 numerical aperture (NA) objective using Zen software (Carl Zeiss Ltd). For live cell imaging of Lact-C2-RFP during autophagosome formation, cells were grown on 35-mm MatTek glass bottomed dishes. Z-stacks were acquired every 20 seconds using a spinning disk confocal microscope, comprising a Nikon Ti-E stand, Nikon 60x 1.45 NA oil immersion lens, Yokogawa CSU-X scanhead, Andor iXon 897 EM-CCD camera and Andor laser combiner. All imaging with live cells was performed within incubation chambers at 37°C and 5% CO_2_. Image acquisition and analysis was performed with Andor iQ3 (Andor Technology, UK) and ImageJ. For Lact-C2-RFP analysis during SMAC, z-stacks were taken using a Confocal Zeiss LSM 780 microscope (Carl Zeiss Ltd) equipped with a 40x oil immersion 1.40 numerical aperture (NA) objective using Zen software (Carl Zeiss Ltd).

For live imaging and quantification of phagocytosis, cells were grown on 35-mm MatTek glass bottomed dishes and z-stacks acquired using a Confocal Zeiss LSM 780 microscope (Carl Zeiss Ltd) equipped with a 40x oil immersion 1.40 numerical aperture (NA) objective using Zen software (Carl Zeiss Ltd). Samples were imaged and maintained in an environment chamber at 37°C and 5% CO_2_. For phagocytosis quantification CellMask (ThermoFisher, C10046) was added to cells prior to imaging. Phagocytosis was quantified as the number of phagosomes per cell that were CellMask negative. Image analysis was performed using Image J software.

#### LC3B-ATG4 complex modelling

The following X-ray crystal structures were used for making models used in illustrations: ATG4B-LC3B 1-120 complex (human), 2z0e.pdb (1.9 Å; Rfree 0.23) ^34^, ATG4B (human), 2cy7.pdb (1.9 Å; Rfree 0.25) ^35^, PE head group, 3PE.pdb from 6tzk.pdb (1.8 Å; Rfree 0.18) ^47^, PS head group, PSF.pdb from 3bib.pdb (2.5 Å; Rfree 0.25) ^48^. The model of LC3B-PE was made by superimposing the ethanolamine moiety of PE (3PE.pdb) on the backbone (Cα), side chain (Cβ) and N of THR121 of LC3B (2z0e.pdb) and an amide bond built between the ethanolamine N and C-terminal carboxyl of GLY120 of LC3B (2z0e.pdb) using PyMOL (The PyMOL Molecular Graphics System, Version 2.2, Schrödinger, LLC). The phosphate of the PE head group was positioned to avoid clashes with ATG4B atoms (2z0e.pdb) by rotation about the ethanolamine carbons. The model of LC3B-PS was made in the same manner using the head group of PS (3PSF.pdb). The ATG4B-LC3B structure contains an HIS280ALA mutation to facilitate stable complex formation. The ALA280 of 2z0e.pdb was changed to a HIS and its position modelled on that found in the native structure of ATG4B (2cy7.pdb) in order to illustrate the active site of ATG4B.

### Statistics

Two-tailed Ratio-paired t-test or student t-test, using Graph Pad, were performed as indicated.

## Acknowledgements

We thank Nick Ktistakis, Len Stephens, Phill Hawkins, Simon Cook and members of the Florey lab for helpful discussions and critical review of the manuscript. This work was supported by grants from the BBSRC, BB/P013384/1 (BBS/E/B/000C0432 and BBS/E/B/000C0434), BB/R019258/1 and Cancer Research UK Career Development award C47718/A16337. BI Mass Spectrometry Facility was supported by a BBSRC Core Capability Grant. This work was partly supported by the Research Council of Norway through its Centres of Excellence funding scheme, project number 262652. We dedicate this work to our friend and colleague Prof. Michael Wakelam, who sadly passed away in March 2020.

## Author contributions

J.D. and O.F. conceived of, designed and carried out experiments and wrote the paper. A.H.L. designed and performed liposome assays. K.S. generated cell lines and carried out experiments (GABARAP and RAW264.7 cell LAP assays). S.R.C. purified ATG4 and RavZ proteins. M.I.W. generated computational models of ATG4B-LC3-II. E.M. and R.U. characterised ATG4D CRISPR KO cell line. A.F.L. and M.J.W. analysed global lipidomics. D.O. and J.W. designed the mass spectrometry experiments and generated data. A.S. and R.B. provided reagents and expertise.

## Data availability

Source Data for Figures 1-4 and Extended Data Figures 2, 3, 5, 6, are provided within the paper. All other data are available from the corresponding author upon request.

